## Supplemental data Dudziak_etal_2025_v2.pdf for "The Spc105/Kre28 complex promotes mitotic error correction by outer kinetochore recruitment of Ipl1/Sli15"

#### Table of contents

**Appendix Figure S1**

**Appendix Figure S1 – Figure Legend**

**Appendix Figure S2**

**Appendix Figure S2 – Figure Legend**

**Appendix Figure S3**

**Appendix Figure S3 – Figure Legend**

**Table S1 – Yeast strains used in the study**

### Appendix Figure S1

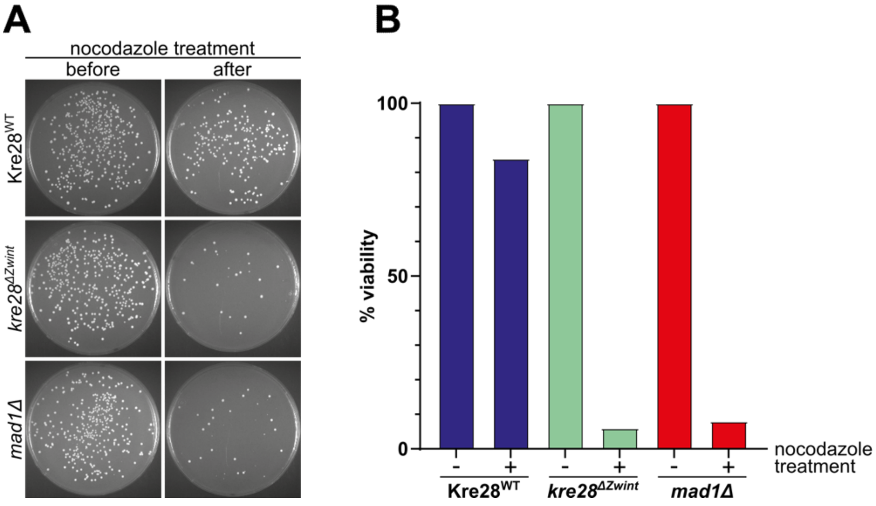

**Appendix Figure S1 – Supplement: Viability assay after nocodazole treatment**
Kre28$^{WT}$, *kre28$^{\Delta Zwint}$* and *mad1Δ* cells were treated with nocodazole for 2 hours. Before and after treatment, a defined number of cells was plated on YEPD plates and incubated at 30 °C. After two days, the number of colonies was counted. A: Images of the plates. B: Quantification of colony formation in the right. For each strain, the number of colonies of the untreated samples was defined as 100 %.

### Appendix Figure S2

Dudziak et al., 2025

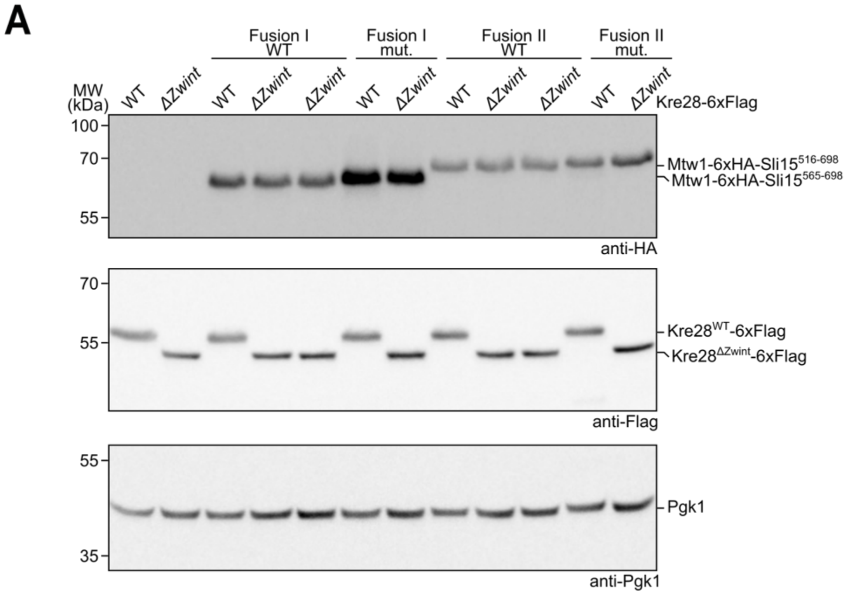

**Appendix Figure S2 – Supplement: Western blot analysis of different Mtw1-Sli15 fusion proteins.** Western blot analysis comparing the expression of Mtw1-6xHA-Sli15 fusion proteins (shorter Fusion I and longer Fusion II) in wild-type form or in mutant form preventing Ipl1 binding. The respective strains were used for serial dilution assays shown in Figure 6B. Pgk1 served as loading control.

### Appendix Figure S3

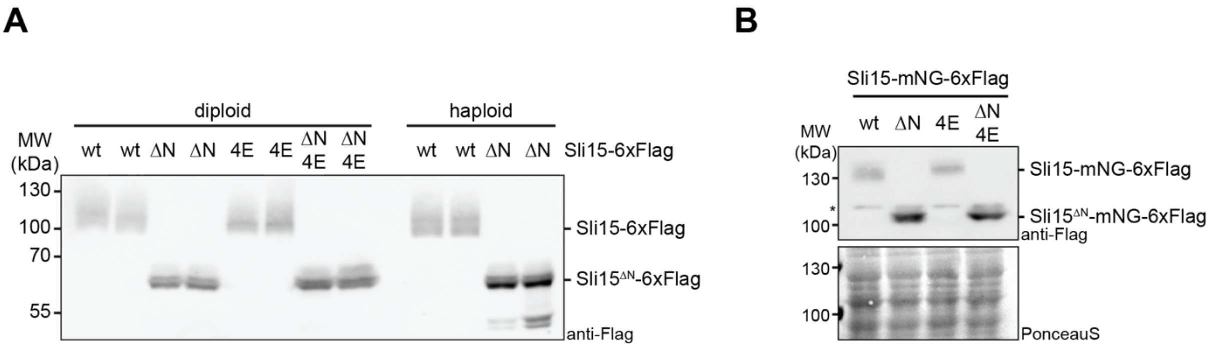

**Appendix Figure S3**. **A**. Western blot analysis of Sli15 replacement constructs in diploid and haploid cells. Yeast extracts were generated from strains with the indicated genotype and blotted against the Flag tag. Note that 4E mutants were not recovered in haploid cells. **B:** Western blot analysis of Sli15-mNeonGreen constructs in diploid cells.

#### Table S1: Yeast strains used in this study

| Name | Genotype | MAT | His | Ura | Lys | Ade | Leu | Figure |
|---|---|---|---|---|---|---|---|---|
| ADY560 | Kre28/Kre28$^{WT}$-6xFlag::Ura3 | a/α | *his3Δ200/ his3Δ200* | | Lys2/*lys2-801* | Ade2/*ade2-1* | *leu2-3,112/leu2-3,112* | Fig 1 |
| ADY561 | Kre28/*kre28*$^{\Delta Zwint}$-6xFlag::Ura3 | a/α | *his3Δ200/ his3Δ200* | | Lys2/*lys2-801* | Ade2/*ade2-1* | *leu2-3,112/leu2-3,112* | Fig 1 |
| ADY784 | Kre28/*kre28*$^{\Delta RWD}$-6xFlag::Ura3 | a/α | *his3Δ200/ his3Δ200* | | Lys2/*lys2-801* | Ade2/*ade2-1* | *leu2-3,112/leu2-3,112* | Fig 1 |
| ADY785 | Kre28/*kre28*$^{\Delta Zwint\ \Delta RWD}$-6xFlag::Ura3 | a/α | *his3Δ200/ his3Δ200* | | Lys2/*lys2-801* | Ade2/*ade2-1* | *leu2-3,112/leu2-3,112* | Fig 1 |
| DDY904 | wildtype strain background | α | *his3Δ200* | | *lys2-801* | | *leu2-3,112* | Fig 1 |
| DDY1502 | *mad1*Δ::His3 | a | | *ura3-52* | | *ade2-101* | *leu2-3,112* | Fig 2 |
| ADY565 | Kre28$^{WT}$-6xFlag::Ura3 | α | *his3Δ200* | | *lys2-801* | | *leu2-3,112* | Fig 1, 6, Fig1 S2 |
| ADY567 | *kre28*$^{\Delta Zwint}$-6xFlag::Ura3 | α | *his3Δ200* | | *lys2-801* | | *leu2-3,112* | Fig 1, 6, Fig1 S2 |
| ADY789 | *kre28*$^{\Delta RWD}$-6xFlag::Ura3 | α | *his3Δ200* | | *lys2-801* | | *leu2-3,112* | Fig 1 |
| JPY25.1 | *kre28*Δ::His3, *ura3-52*::Kre28$^{WT}$-6xFlag::Ura3 | a | | | | *ade2-1* | *leu2-3,112* | Fig EV1 |
| FHY21a | *kre28*Δ::His3, *ura3-52*::*kre28*$^{\Delta Zwint}$-6xFlag::Ura3 | a | | | | *ade2-1* | *leu2-3,112* | Fig EV1 |
| JPY21.3 | *kre28*Δ::His3, *ura3-52*::*kre28*$^{\Delta 91\text{-}102}$-6xFlag::Ura3 | a | | | | *ade2-1* | *leu2-3,112* | Fig EV1 |
| JPY21.4 | *kre28*Δ::His3, *ura3-52*::*kre28*$^{\Delta 91\text{-}102}$-6xFlag::Ura3 | a | | | | *ade2-1* | *leu2-3,112* | Fig EV1 |
| JPY22.2 | *kre28*Δ::His3, *ura3-52*::*kre28*$^{\Delta 77\text{-}90}$-6xFlag::Ura3 | α | | | | *ade2-1* | *leu2-3,112* | Fig EV1 |
| JPY22.4 | *kre28*Δ::His3, *ura3-52*::*kre28*$^{\Delta 77\text{-}90}$-6xFlag::Ura3 | a | | | | | *leu2-3,112* | Fig EV1 |
| JPY23.3 | *kre28*Δ::His3, *ura3-52*::*kre28*$^{\Delta 103\text{-}113}$-6xFlag::Ura3 | α | | | | *ade2-1* | *leu2-3,112* | Fig EV1 |
| JPY23.4 | *kre28*Δ::His3, *ura3-52*::*kre28*$^{\Delta 103\text{-}113}$-6xFlag::Ura3 | a | | | *lys2-801* | *ade2-1* | *leu2-3,112* | Fig EV1 |
| FHY19b | *kre28*Δ::His3, *ura3-52*::Kre28$^{WT}$-6xFlag::Ura3 | α | | | *lys2-801* | | *leu2-3,112* | Fig EV1 |
| FHY22b | *kre28*Δ::His3, *ura3-52*::*kre28*$^{\Delta Zwint}$-6xFlag::Ura3 | α | | | *lys2-801* | | *leu2-3,112* | Fig EV1 |
| ADY623.1 | *kre28*Δ::His3, *ura3-52*::*kre28*$^{4A}$-6xFlag::Ura3 (L95A, K96A, Y99A, E101A) | α | | | | | *leu2-3,112* | Fig EV1 |
| ADY623.2 | *kre28*Δ::His3, *ura3-52*::*kre28*$^{4A}$-6xFlag::Ura3 (L95A, K96A, Y99A, E101A) | α | | | | | *leu2-3,112* | Fig EV1 |
| ADY625.1 | *kre28*Δ::His3, *ura3-52*::*kre28*$^{3A}$-6xFlag::Ura3 (E103A, L105A, D106A) | α | | | | | *leu2-3,112* | Fig EV1 |
| ADY625.2 | *kre28*Δ::His3, *ura3-52*::*kre28*$^{3A}$-6xFlag::Ura3 (E103A, L105A, D106A) | α | | | *lys2-801* | | *leu2-3,112* | Fig EV1 |
| ADY627.1 | *kre28*Δ::His3, *ura3-52*::*kre28*$^{6A}$-6xFlag::Ura3 (F108A, F109A, R110A, F111A, T112A, L113A) | α | | | | | *leu2-3,112* | Fig EV1 |
| ADY627.2 | *kre28*Δ::His3, *ura3-52*::*kre28*$^{6A}$-6xFlag::Ura3 (F108A, F109A, R110A, F111A, T112A, L113A) | α | | | | *ade2-1* | *leu2-3,112* | Fig EV1 |
| ADY573 | Kre28$^{WT}$-6xFlag::Ura3, Pds1-18xMyc::Leu2 | a | *his3Δ200* | | | *ade2-1* | | Fig 2 |
| ADY574 | *kre28*$^{\Delta Zwint}$-6xFlag::Ura3, Pds1-18xMyc::Leu2 | a | *his3Δ200* | | | *ade2-1* | | Fig 2 |
| ADY539 | *mad1*Δ::His3, Pds1-18xMyc::Leu2 | a | | *ura3-52* | | *ade2-1* | | Fig 2 |
| DDY902 | wildtype strain background | a | *his3Δ200* | *ura3-52* | | *ade2-1* | *leu2-3,112* | Fig 2 |
| ADY564 | Kre28$^{WT}$-6xFlag::Ura3 | a | *his3Δ200* | | | *ade2-1* | *leu2-3,112* | Fig 2 |
| ADY566 | *kre28*$^{\Delta Zwint}$-6xFlag::Ura3 | a | *his3Δ200* | | | *ade2-1* | *leu2-3,112* | Fig 2 |
| ADY581.1 | Kre28$^{WT}$-6xFlag::Ura3, *mad1*Δ::His3 | a | | | | *ade2-1* | | Fig 2 |

| | | | | | | | | |
|---|---|---|---|---|---|---|---|---|
| ADY581.2 | Kre28$^{WT}$-6xFlag::Ura3, *mad1*Δ::His3 | a | | | *lys2-801* | *ade2-1* | | Fig 2 |
| ADY583.1/2 | *kre28*$^{\Delta Zwint}$-6xFlag::Ura3, *mad1*Δ::His3 | a | | | | *ade2-1* | | Fig 2 |
| ADY562 | Kre28$^{WT}$-6xFlag::Ura3, *cdc15-2*::Leu2, *his3Δ200*::pCup1-LacI-GFP::His3, CEN XII-prox.::LacO::Leu2, Spc42-mCherry::KanMx6 | a | | | | *ade2-1* | | Fig 2, 4 |
| ADY563 | *kre28*$^{\Delta Zwint}$-6xFlag::Ura3, *cdc15-2*::Leu2, *his3Δ200*::pCup1-LacI-GFP::His3, CEN XII-prox.::LacO::Leu2, Spc42-mCherry::KanMx6 | a | | | | *ade2-1* | | Fig 2, 4 |
| ADY620 | Kre28/Kre28$^{WT}$-6xFlag::Ura3, Ipl1/*ipl1-1* | a/α | *his3Δ200/his3Δ200* | | Lys2/*lys2-801* | Ade2/*ade2-1* | *leu2-3,112/leu2-3,112* | Fig 2 |
| ADY621 | Kre28/*kre28*$^{\Delta Zwint}$-6xFlag::Ura3, Ipl1/*ipl1-1* | a/α | *his3Δ200/his3Δ200* | | Lys2/*lys2-801* | Ade2/*ade2-1* | *leu2-3,112/leu2-3,112* | Fig 2 |
| FHY18a | *kre28*Δ::His3, *ura3-52*::Kre28$^{WT}$-6xFlag::Ura3 | a | | | | *ade2-1* | *leu2-3,112* | Fig 2 |
| SWY2265 | *mad1Δ*::KanMx6 | α | *his3Δ200* | *ura3-52* | *lys2-801* | | *leu2-3,112* | Fig2 |
| ADY661 | Kre28$^{WT}$-6xFlag::Ura3, Mtw1-GFP::KanMx6, Spc42-RFP::His3 | a | | | | | *leu2-3,112* | Fig 3 |
| ADY663 | *kre28*$^{\Delta Zwint}$-6xFlag::Ura3, Mtw1-GFP::KanMx6, Spc42-RFP::His3 | a | | | | | *leu2-3,112* | Fig 3 |
| ADY674 | Kre28$^{WT}$-6xFlag::Ura3, Nuf2-GFP::KanMx6, Spc42-RFP::His3 | a | | | | | *leu2-3,112* | Fig 3 |
| ADY668 | *kre28*$^{\Delta Zwint}$-6xFlag::Ura3, Nuf2-GFP::KanMx6, Spc42-RFP::His3 | a | | | | | *leu2-3,112* | Fig 3 |
| ADY638 | Kre28$^{WT}$-6xFlag::Ura3, Spc105-GFP::KanMx6, Spc42-RFP::His3 | a | | | | | *leu2-3,112* | Fig 3 |
| ADY641 | *kre28*$^{\Delta Zwint}$-6xFlag::Ura3, Spc105-GFP::KanMx6, Spc42-RFP::His3 | a | | | *lys2-801* | | *leu2-3,112* | Fig 3 |
| ADY791 | *kre28*Δ::His3, *ura3-52*::GFP-Kre28$^{WT}$-6xFlag::Ura3, Spc42-mCherry::KanMx6 | α | | | *lys2-801* | | *leu2-3,112* | Fig EV2 |
| ADY793 | *kre28*Δ::His3, *ura3-52*::GFP-*kre28*$^{\Delta Zwint}$-6xFlag::Ura3, Spc42-mCherry::KanMx6 | α | | | *lys2-801* | | *leu2-3,112* | Fig EV2 |
| ADY639 | Kre28$^{WT}$-6xFlag::Ura3, Spc105-GFP::KanMx6, Spc42-RFP::His3 | α | | | *lys2-801* | | *leu2-3,112* | Fig EV2 |
| ADY642 | *kre28*$^{\Delta Zwint}$-6xFlag::Ura3, Spc105-GFP::KanMx6, Spc42-RFP::His3 | α | | | *lys2-801* | | *leu2-3,112* | Fig EV2 |
| ADY662 | Kre28$^{WT}$-6xFlag::Ura3, Mtw1-GFP::KanMx6, Spc42-RFP::His3 | α | | | *lys2-801* | | *leu2-3,112* | Fig EV2 |
| ADY664 | *kre28*$^{\Delta Zwint}$-6xFlag::Ura3, Mtw1-GFP::KanMx6, Spc42-RFP::His3 | α | | | *lys2-801* | | *leu2-3,112* | Fig EV2 |
| ADY675 | Kre28$^{WT}$-6xFlag::Ura3, Nuf2-GFP::KanMx6, Spc42-RFP::His3 | α | | | | | *leu2-3,112* | Fig EV2 |
| ADY669 | *kre28*$^{\Delta Zwint}$-6xFlag::Ura3, Nuf2-GFP::KanMx6, Spc42-RFP::His3 | α | | | | | *leu2-3,112* | Fig EV2 |
| ADY790 | *kre28*Δ::His3, *ura3-52*::GFP-Kre28$^{WT}$-6xFlag::Ura3, Spc42-mCherry::KanMx6 | a | | | *lys2-801* | | *leu2-3,112* | Fig EV2 |
| ADY792 | *kre28*Δ::His3, *ura3-52*::GFP-*kre28*$^{\Delta Zwint}$-6xFlag::Ura3, Spc42-mCherry::KanMx6 | a | | | *lys2-801* | | *leu2-3,112* | Fig EV2 |
| ADY712 | Kre28$^{WT}$-GFP::Ura3, Spc42-RFP::His3 | a | | | | | *leu2-3,112* | Fig EV2 |
| ADY713 | Kre28$^{WT}$-GFP::Ura3, Spc42-RFP::His3 | α | | | *lys2-801* | | *leu2-3,112* | Fig EV2 |
| ADY714 | *kre28*$^{\Delta Zwint}$-GFP::Ura3, Spc42-RFP::His3 | a | | | | | *leu2-3,112* | Fig EV2 |
| ADY715 | *kre28*$^{\Delta Zwint}$-GFP::Ura3, Spc42-RFP::His3 | α | | | *lys2-801* | | *leu2-3,112* | Fig EV2 |
| ADY654 | Kre28$^{WT}$-6xFlag::Ura3, Sgo1-GFP::KanMx6, Spc42-RFP::His3 | a | | | | | *leu2-3,112* | Fig EV2 |
| ADY656 | *kre28*$^{\Delta Zwint}$-6xFlag::Ura3, Sgo1-GFP::KanMx6, Spc42-RFP::His3 | a | | | | | *leu2-3,112* | Fig EV2 |
| ADY697 | Kre28/Kre28$^{WT}$-6xFlag::Ura3, Sgo1/*sgo1Δ*::natNT2 | a/α | *his3Δ200/his3Δ200* | | Lys2/*lys2-801* | Ade2/*ade2-1* | *leu2-3,112/leu2-3,112* | Fig 2 |
| ADY698 | Kre28/*kre28*$^{\Delta Zwint}$-6xFlag::Ura3, Sgo1/*sgo1Δ*::natNT2 | a/α | *his3Δ200/his3Δ200* | | Lys2/*lys2-801* | Ade2/*ade2-1* | *leu2-3,112/leu2-3,112* | Fig 2 |

| | | | | | | | | |
|---|---|---|---|---|---|---|---|---|
| ADY707 | Kre28/Kre28$^{WT}$-6xFlag::Ura3, Sli15/sli15-3 | a/α | *his3Δ200/ his3Δ200* | | Lys2/*lys2-801* | Ade2/*ade2-1* | *leu2-3,112/leu2-3,112* | Fig 2 |
| ADY708 | Kre28/*kre28*$^{\Delta Zwint}$-6xFlag::Ura3, Sli15/*sli15-3* | a/α | *his3Δ200/ his3Δ200* | | Lys2/*lys2-801* | Ade2/*ade2-1* | *leu2-3,112/leu2-3,112* | Fig 2 |
| ADY618 | Kre28/Kre28$^{WT}$-6xFlag::Ura3, Ndc80/*ndc80-1* | a/α | *his3Δ200/ his3Δ200* | | Lys2/*lys2-801* | Ade2/*ade2-1* | *leu2-3,112/leu2-3,112* | Fig 2 |
| ADY619 | Kre28/*kre28*$^{\Delta Zwint}$-6xFlag::Ura3, Ndc80/*ndc80-1* | a/α | *his3Δ200/ his3Δ200* | | Lys2/*lys2-801* | Ade2/*ade2-1* | *leu2-3,112/leu2-3,112* | Fig 2 |
| ADY775 | Kre28/Kre28$^{WT}$-6xFlag::Ura3, Ctf19/Ctf19$^{WT}$-13xMyc::His3 | a/α | | | Lys2/*lys2-801* | Ade2/*ade2-1* | *leu2-3,112/leu2-3,112* | Fig 2 |
| ADY776 | Kre28/Kre28$^{WT}$-6xFlag::Ura3, Ctf19/*ctf19*$^{\Delta C\text{-}RWD\ (270\text{-}369)}$-13xMyc::His3 | a/α | | | Lys2/*lys2-801* | Ade2/*ade2-1* | *leu2-3,112/leu2-3,112* | Fig 2 |
| ADY777 | Kre28/*kre28*$^{\Delta Zwint}$-6xFlag::Ura3, Ctf19/Ctf19$^{WT}$-13xMyc::His3 | a/α | | | Lys2/*lys2-801* | Ade2/*ade2-1* | *leu2-3,112/leu2-3,112* | Fig 2 |
| ADY778 | Kre28/*kre28*$^{\Delta Zwint}$-6xFlag::Ura3, Ctf19/*ctf19*$^{\Delta C\text{-}RWD\ (270\text{-}369)}$-13xMyc::His3 | a/α | | | Lys2/*lys2-801* | Ade2/*ade2-1* | *leu2-3,112/leu2-3,112* | Fig 2 |
| ADY616 | Kre28/Kre28$^{WT}$-6xFlag::Ura3, Dam1/*dam1-1*::KanMx6 | a/α | *his3Δ200/ his3Δ200* | | Lys2/*lys2-801* | Ade2/*ade2-1* | *leu2-3,112/leu2-3,112* | Fig 2 |
| ADY617 | Kre28/*kre28*$^{\Delta Zwint}$-6xFlag::Ura3, Dam1/*dam1-1*::KanMx6 | a/α | *his3Δ200/ his3Δ200* | | Lys2/*lys2-801* | Ade2/*ade2-1* | *leu2-3,112/leu2-3,112* | Fig 2 |
| ADY315 | *duo1*$^{\Delta SxIP}$::Leu2 | a | *his3Δ200* | *ura3-52* | | *ade2-1* | | Fig 2 |
| ADY577.1/2 | Kre28$^{WT}$-6xFlag::Ura3, *duo1*$^{\Delta SxIP}$ | a | *his3Δ200* | | | *ade2-1* | | Fig 2 |
| ADY579.1 | Kre28$^{\Delta Zwint}$-6xFlag::Ura3, *duo1*$^{\Delta SxIP}$ | a | *his3Δ200* | | | *ade2-1* | | Fig 2 |
| ADY579.2 | Kre28$^{\Delta Zwint}$-6xFlag::Ura3, *duo1*$^{\Delta SxIP}$ | a | *his3Δ200* | | *lys2-801* | *ade2-1* | | Fig 2 |
| SWY389A | *cnn1Δ*::His3 | a | | *ura3-52* | | *ade2-1* | *leu2-3,112* | Fig 2 |
| ADY685 | Kre28$^{WT}$-6xFlag::Ura3, *cnn1Δ*::His3 | a | | | | *ade2-1* | *leu2-3,112* | Fig 2 |
| ADY686 | Kre28$^{WT}$-6xFlag::Ura3, *cnn1Δ*::His3 | α | | | *lys2-801* | | *leu2-3,112* | Fig 2 |
| ADY687 | *kre28*$^{\Delta SxIP}$-6xFlag::Ura3, *cnn1Δ*::His3 | a | | | | *ade2-1* | *leu2-3,112* | Fig 2 |
| ADY688 | *kre28*$^{\Delta SxIP}$-6xFlag::Ura3, *cnn1Δ*::His3 | α | | | *lys2-801* | | *leu2-3,112* | Fig 2 |
| ADY768 | Kre28$^{WT}$-6xFlag::Ura3, *lys2-801*::Mtw1-6xHA-Sli15$^{565\text{-}698\ (WT)}$::Lys2 | α | *his3Δ200* | | | | *leu2-3,112* | Fig 4 |
| ADY769 | *kre28*$^{\Delta Zwint}$-6xFlag::Ura3, *lys2-801*::Mtw1-6xHA-Sli15$^{565\text{-}698\ (WT)}$::Lys2 | α | *his3Δ200* | | | | *leu2-3,112* | Fig 4 |
| ADY802 | Kre28$^{WT}$-6xFlag::Ura3, *lys2-801*::Mtw1-6xHA-Sli15$^{516\text{-}698\ (WT)}$::Lys2 | α | *his3Δ200* | | | | *leu2-3,112* | Fig 4 |
| ADY803 | *kre28*$^{\Delta Zwint}$-6xFlag::Ura3, *lys2-801*::Mtw1-6xHA-Sli15$^{516\text{-}698\ (WT)}$::Lys2 | α | *his3Δ200* | | | | *leu2-3,112* | Fig 4 |
| ADY794 | Kre28$^{WT}$-6xFlag::Ura3, *lys2-801*::Mtw1-6xHA-Sli15$^{565\text{-}698\ (W646G,\ F680A)}$::Lys2 | α | *his3Δ200* | | | | *leu2-3,112* | Fig 4 |
| ADY795 | *kre28*$^{\Delta Zwint}$-6xFlag::Ura3, *lys2-801*::Mtw1-6xHA-Sli15$^{565\text{-}698\ (W646G,\ F680A)}$::Lys2 | α | *his3Δ200* | | | | *leu2-3,112* | Fig 4 |
| ADY840 | Kre28$^{WT}$-6xFlag::Ura3, *lys2-801*::Mtw1-6xHA-Sli15$^{516\text{-}698\ (W646G,\ F680A)}$::Lys2 | α | *his3Δ200* | | | | *leu2-3,112* | Fig 4 |
| ADY841 | *kre28*$^{\Delta Zwint}$-6xFlag::Ura3, *lys2-801*::Mtw1-6xHA-Sli15$^{516\text{-}698\ (W646G,\ F680A)}$::Lys2 | α | *his3Δ200* | | | | *leu2-3,112* | Fig 4 |
| ADY836 | Kre28$^{WT}$-6xFlag::Ura3, *cdc15-2*::Leu2, *his3Δ200*::pCup1-LacI-GFP::His3, CEN XII-prox.::LacO::Leu2, Spc42-mCherry::KanMx6, *lys2-801*::Mtw1-6xHA-Sli15$^{565\text{-}698\ (WT)}$::natNT2::Lys2 | α | | | | *ade2-1* | | Fig 4 |
| ADY837 | *kre28*$^{\Delta Zwint}$-6xFlag::Ura3, *cdc15-2*::Leu2, *his3Δ200*::pCup1-LacI-GFP::His3, CEN XII-prox.::LacO::Leu2, Spc42-mCherry::KanMx6, *lys2-801*::Mtw1-6xHA-Sli15$^{565\text{-}698\ (WT)}$::natNT2::Lys2 | α | | | | *ade2-1* | | Fig 4 |

| | | | | | | | | |
|---|---|---|---|---|---|---|---|---|
| ADY838 | Kre28$^{WT}$-6xFlag::Ura3, *cdc15-2*::Leu2, *his3Δ200*::pCup1-LacI-GFP::His3, CEN XII-prox.::LacO::Leu2, Spc42-mCherry::KanMx6, *lys2-801*::Mtw1-6xHA-Sli15$^{516\text{-}698\ (WT)}$::natNT2::Lys2 | α | | | | *ade2-1* | | Fig 4 |
| ADY839 | *kre28*$^{\Delta Zwint}$-6xFlag::Ura3, *cdc15-2*::Leu2, *his3Δ200*::pCup1-LacI-GFP::His3, CEN XII-prox.::LacO::Leu2, Spc42-mCherry::KanMx6, *lys2-801*::Mtw1-6xHA-Sli15$^{516\text{-}698\ (WT)}$::natNT2::Lys2 | α | | | | *ade2-1* | | Fig 4 |
| ADY816 | Kre28 WT-6xFlag, Spc105-TurboID-3xMyc::KanMx6 | a | *his3Δ200* | | | *ade2-1* | *leu2-3,112* | Fig 5 |
| ADY828 | Spc105-TurboID-3xMyc::KanMx6, Sli15 WT-6xFlag::His3 | a | | *ura3-52* | | *ade2-1* | *leu2-3,112* | Fig 5 |
| STY327 | Cin8-6xFlag::URA3 | a | *his3Δ200* | | | | *leu2-3,112* | Fig 5 |
| STY328 | Cin8-6xFlag::URA3, Spc105-TurboID-3xMyc::KanMx6 | a | *his3Δ200* | | | | *leu2-3,112* | Fig 5 |
| ADY865 | sli15::Sli15-6xFlag::His3 | a | | | | | | Fig 7 |
| ADY866 | sli15::Sli15(delta2-230)-6xFlag::His3 | a | | | | | | Fig 7 |
| ADY873 | Mtw1/Mtw1-mCherry::KanMx6, Sli15/Sli15-mNeonGreen-6xFlag | a/α | | | | | | Fig 7 |
| ADY874 | Mtw1/Mtw1-mCherry::KanMx6, Sli15/Sli15(delta2-230)-mNeonGreen-6xFlag | a/α | | | | | | Fig 7 |
| ADY875 | Mtw1/Mtw1-mCherry::KanMx6, Sli15/Sli15(R231E R232E L239E K242E)-mNeonGreen-6xFlag | a/α | | | | | | Fig 7 |
| ADY876 | Mtw1/Mtw1-mCherry::KanMx6, Sli15/Sli15(delta2-230, R231E R232E L239E K242E)-mNeonGreen-6xFlag | a/α | | | | | | Fig 7 |
| ADY877 | Sli15-aid*-9xmyc::KanMX6 ura3-52::pTEF1-OsTIR1_tADH1::URA3 | a | *his3Δ200* | | | | | Fig EV5 |
| ADY878 | Sli15-aid::KanMX with Sli15-wt-6xFlag::LEU2 | a | *his3Δ200* | | | | | Fig EV5 |
| ADY879 | Sli15-aid::KanMX with Sli15-deltaN-6xFlag::LEU | a | *his3Δ200* | | | | | Fig EV5 |
| ADY880 | Sli15-aid::KanMX with Sli15-(R231E R232E L239E K242E)-6xFlag::LEU2 | a | *his3Δ200* | | | | | Fig EV5 |
| ADY881 | Sli15-aid::KanMX with Sli15(delta2-230, R231E R232E L239E K242E)-6xFlag | a | *his3Δ200* | | | | | Fig EV5 |
| ADY882 | Sli15-aid::KanMX with Sli15-2E (R231E R1232E)-6xFlag::LEU2 | a | *his3Δ200* | | | | | Fig EV5 |
| ADY883 | Sli15-aid::KanMX with Sli15-1E(K242E)-6xFlag::LEU2 | a | *his3Δ200* | | | | | Fig EV5 |
| ADY884 | Spc105-FRB::KanMX, ura3-52::Spc105-wt::URA3 | a | | | | | | Fig EV5 |
| ADY885 | Spc105-FRB::KanMX, ura3-52::Spc105 E578K E596K::URA3 | a | | | | | | Fig EV5 |
| ADY886 | Spc105-FRB::KanMX, ra3-52::Spc105 E510K E578K E596K::URA3 | a | | | | | | Fig EV5 |
